# The p48 isoform of the PA2G4/EBP1/ITAF45 oncoprotein is required for the encephalomyocarditis virus IRES-driven translation initiation

**DOI:** 10.1101/2025.08.22.671869

**Authors:** Artem S. Kushchenko, Violetta A. Golovko, Eugenia A. Panova, Anastasia P. Sukhunina, Ekaterina E. Gladneva, Alexandr Y. Krasota, Yury Y. Ivin, Anastasia V. Poteryakhina, Vadim I. Agol, Sergey E. Dmitriev

**Affiliations:** Belozersky Institute of Physico-Chemical Biology, Lomonosov Moscow State University, Moscow, 119234, Russia; Faculty of Bioengineering and Bioinformatics, Lomonosov Moscow State University, Moscow, 119234, Russia; Biological Faculty, Lomonosov Moscow State University, Moscow, 119234, Russia; Chumakov Federal Scientific Center for Research and Development of Immune-and-Biological Products of Russian Academy of Sciences (Institute of Poliomyelitis), Moscow, 108819, Russia; Engelhardt Institute of Molecular Biology, Russian Academy of Sciences, 119991 Moscow, Russia

**Keywords:** internal ribosome entry site, IRES trans-acting factor ITAF45, proliferation-associated protein 2G4, p48 proteoform, ribosome biogenesis factor EBP1, Mengovirus, Mengo IRES, EMCV IRES, FMDV IRES, CRISPR screening, mRNA transfection, replicon

## Abstract

Internal ribosome entry sites (IRESs) enable cap-independent initiation of picornaviral RNA translation and, together with canonical translation initiation factors, typically require specific cellular proteins known as IRES *trans*-acting factors (ITAFs). While the type II IRES of foot-and-mouth disease virus (FMDV, an aphthovirus) has been shown to depend on the oncoprotein ITAF45, also known as Proliferation-Associated 2G4 (PA2G4) or ErbB-3 receptor Binding Protein (EBP1), for *in vitro* assembly of the 48S pre-initiation complex, some related type II IRESs, such as that of encephalomyocarditis virus (EMCV, a cardiovirus), can form the initiation complex independently of ITAF45. In this study, we performed a genome-wide CRISPR screen and identified knockouts of *PA2G4*/*EBP1*/*ITAF45* in cells that survive EMCV infection, suggesting an important role for this factor. We show that the p48 isoform of ITAF45, but not the p42 isoform, is crucial for efficient EMCV/Mengovirus replication and for propagation of replicons in human cell culture. Loss of ITAF45 markedly diminishes EMCV and FMDV IRES activities, which can be rescued by re-expression of ITAF45-p48. Interestingly, cell-free translation assays reveal that EMCV IRES activity is less ITAF45-dependent *in vitro*, in contrast to FMDV, raising questions about the versatile functions of ITAFs in IRES-driven translation. These findings reveal an isoform-specific function of ITAF45 in supporting cardiovirus infection and provide new insights into the complex regulation of IRES-driven translation, with implications for developing targeted antiviral strategies.

## INTRODUCTION

Viruses are obligate intracellular parasites that exploit the host translational machinery to produce their own proteins (1). Picornaviruses represent a diverse group of small non-enveloped positive-strand RNA viruses whose entire life cycle occurs in the cytoplasm (2-4). The genomic RNA of picornaviruses encodes a single polyprotein, which is then proteolytically processed into several viral proteins (5). Encephalomyocarditis virus (EMCV) serves as a model representative of cardioviruses, which belong to the *Caphthovirinae* subfamily of picornaviruses (4). First isolated in 1945 (6), EMCV has been established as the causative agent of myocarditis and encephalitis in various animal species (7). EMCV encompasses several subspecies, including Mengovirus (hereafter referred to as Mengo).

The EMCV genomic RNA is 7.8 kb long, has 5’ and 3’ untranslated regions (UTRs) of approximately 800-1,200 nt and 120 nt, respectively; at the 5’ end, it is covalently linked to a 20-amino-acid viral protein, VPg (7,8). The 5’ proximal region of the EMCV 5’ UTR adopts a cloverleaf structure that plays a critical role in viral genome replication, followed by a long poly(C)-tract that affects the virulence, pathogenicity, and tropism of the virus (7,9). The remaining major portion of the 5’ UTR harbors a type II internal ribosome entry site (IRES) (10), a highly structured RNA element approximately 500 nt in length that facilitates ribosome recruitment through direct binding of initiation factors, thereby initiating viral polypeptide synthesis (11-14).

Type II IRESs comprise several RNA hairpins combined into structural domains (numbered as II-V/VI or, according to another nomenclature, H-K/L) (15). The J-K domain specifically binds translation initiation factor (eIF) eIF4G, making the viral mRNA translation independent of cap and eIF4E (16). This domain alone provides cap-independence to an mRNA but it is insufficient for IRES activity, i.e. for the ribosome recruitment into internal parts of a transcript (see (17) and references therein). EMCV IRES ends with an oligopyrimidine tract (Y_n_) followed by a short spacer and three AUG codons, two of which (11^th^ and 12^th^ AUGs) are in the same reading frame and can initiate synthesis of the viral polyprotein. The 11^th^ AUG (located at the position 834 of the 5’ UTR) is utilized as the major start codon (18). In the course of infection, EMCV security protein 2A activates 4E-BP1, which interacts with the cap-binding factor eIF4E (19). This inhibits cap-dependent translation of cellular mRNAs and provides the translational priority to the virus (5,20).

Foot-and-mouth disease virus (FMDV) is a model member of aphthoviruses, another genus of *Caphthovirinae* (2,4), and is a well-known causative agent of foot-and-mouth disease in cattle (21,22). The FMDV genomic RNA (gRNA) also possesses a type II IRES with an overall architecture similar to that of EMCV. Experimental evidence indicates that the structural domains of the FMDV IRES are involved in tertiary interactions impacting its cap-independent activity (23). Notably, FMDV also has two in-frame AUG codons in the initiation region (hereafter called AUG1 and AUG2), but in this case they are separated by 84 nt and the start of polyprotein synthesis occurs predominantly at the second AUG (see (24) and references therein). The delivery of the 43S pre-initiation complex to AUG2 requires scanning that is facilitated by eIF1 (25). Furthermore, FMDV also disrupts the canonical translation initiation mechanism through its L protease, which cleaves eIF4G, thereby suppressing the cap-dependent translation initiation exploited by cellular mRNAs (26). Nevertheless, the C-terminal fragment of eIF4G retains the ability to bind to the J-K domain of the FMDV IRES, facilitating ribosome recruitment to the viral RNA thus sustaining its translation (26).

Type II IRESs utilize canonical eIFs to assemble the 48S pre-initiation complex and position the ribosome at the start codon (25,27,28). However, in addition, these IRESs require cellular RNA-binding proteins, typically not associated with the translation initiation process, known as IRES *trans*-acting factors (ITAFs) (for review, see (11,29)). The prevailing hypothesis posits that ITAFs enhance the stability of the optimal IRES conformation, thereby acting as RNA chaperones and facilitating efficient ribosomal recruitment (30-32). The requirement of viral IRESs for additional cellular factors is particularly intriguing from a medical perspective, as the presence or absence of ITAFs in cells may influence the neurotropism of the virus (33).

The first ITAF identified was PTB/PTBP1 (Polypyrimidine-Tract-Binding Protein), which was found to bind to the IRESs of both EMCV (34-36) and FMDV (37). In uninfected cells, PTB plays a crucial role in alternative pre-mRNA splicing (38). However, PTB has been shown to be either necessary or stimulatory for IRES-directed translation of both EMCV and FMDV (35,39-41), as well as for the assembly of 48S complexes on type II IRESs (27,28,40,42). The extent of stimulation varies depending on the IRES sequence, the purity of the factor preparations, and likely other conditions (39,40,42).

Another ITAF reported to operate on the FMDV IRES is ITAF45 (28). ITAF45, also known as PA2G4 (Proliferation-Associated 2G4) or Ebp1 (ErbB-3 receptor Binding Protein), was initially identified as a multifunctional protein that regulates cell proliferation (43-46). Two isoforms of ITAF45, p42 and p48, differ by 54 amino acid residues at the N-terminus and have distinct interacting partners (43,47). The p42 isoform lacks the first α-helix and nearly a half of the second α-helix present in p48, resulting in a surface enriched with hydrophobic amino acid residues (48). The expression of the pro-proliferative p48 isoform has been detected at a high level in numerous cancer cell types (for review, see (43,44)), whereas the level of the p42 isoform is significantly reduced in cancer cells, where it is selectively ubiquitinated and subsequently degraded (49). The p42 isoform primarily resides in the cytoplasm, whereas the p48 isoform is present in both the cytoplasm and the nucleus (50,51). The nuclear-localized portion of the p48 isoform is found in the nucleolus, presumably participating in ribosome biogenesis (51,52).

ITAF45 is orthologous to the yeast 60S maturation and nuclear export factor Arx1p (53-57). However, mammalian ITAF45 lacks the regions present in Arx1p that are responsible for its interactions with the ribosome maturation factors Alb1p and Rei1p, and it does not interact with nucleoporins, calling into question its complete functional identity to yeast Arx1p (53-55,58-60). Instead, ITAF45 has been found to stably bind to mature 80S ribosomes in mammals (60).

ITAF45/Arx1p exhibits structural homology with proteins belonging to the methionine aminopeptidase (MetAP) family; however, it lacks the corresponding catalytic activity and possesses an additional RNA-binding site at the C-terminus (61,62). ITAF45 and Arx1p bind to the ribosome within the inner ring of the ribosomal polypeptide tunnel (53,54,59,63-65), a region also occupied by catalytically active MetAP (66). In this area, ITAF45 interacts with the uL23 ribosomal protein, a general docking site for several nascent chain-interacting factors (such as chaperons, enzymes, and targeting factors), and 5.8S rRNA (59,63). Additionally, ITAF45 forms extensive contacts with 28S rRNA, particularly with helices H24, H47, H53, and H59, and notably with the expansion segment 27L (ES27L). As a result, ITAF45 is “sandwiched” between the ribosomal tunnel and this expansion segment, simultaneously interacting with several ribosomal RNA and protein elements.

While yeast Arx1p is believed to accompany the pre-60S particles only during the late stages of its biogenesis, mammalian ITAF45 is thought to bind to the ribosomes engaged in active translation, particularly from the initiation stage until the growing peptide chain extends beyond the peptide tunnel (65). Following this phase, ITAF45 dissociates, allowing translation to proceed. For secretory, membrane, or mitochondrial proteins carrying specific localization signals, ITAF45 is replaced by corresponding factors that facilitate the co-translational targeting of the ribosomal complex to the organelle membrane translocon (65).

ITAF45 was also shown to interact with PKR protein kinase, suppress eIF2α phosphorylation (67), and associate with some coding and non-coding RNAs in both the cytoplasm and nucleus (52,68).

In a study by Pilipenko et al., ITAF45 was found to be critical for the reconstitution of the pre-initiation 48S complex on the FMDV IRES, along with PTB and a subset of canonical initiation factors (28). The absence of either of these ITAFs resulted in inefficient assembly of the 48S complex at both AUG1 and AUG2 codons (25,28). The binding of ITAF45 to the FMDV IRES was independently confirmed using a riboproteomic approach (69). In contrast, the subset of canonical initiation factors was found to be sufficient for the assembly of the 48S complex on the EMCV IRES (27). While the addition of PTB significantly enhanced the efficiency of complex formation and was essential under certain conditions (27,42), the inclusion of ITAF45 was claimed to have no effect on the 48S complex assembly (28). These findings were further confirmed by ITAF45 knockdown experiments in cultured cells, which demonstrated a reduction to half the efficiency of FMDV IRES activity (62,70), while the activity of the EMCV IRES remained unaffected (62).

However, direct RNA-protein interaction assays revealed that ITAF45 binds to both FMDV and EMCV IRESs (28,30,62), thereby raising questions regarding its potential involvement in the activity of the EMCV IRES as well. In this study, we identify ITAF45 as a host factor essential for the EMCV/Mengo virus life cycle in cultured human cells. We demonstrate that disruption of the ITAF45 gene renders cells resistant to infection, significantly reducing viral yield and RNA replication by inhibiting IRES-dependent translation. Moreover, expression of the p48 (but not p42) isoform rescues translation in cultured cells, as well as viral propagation. These findings highlight a specific functional role for the p48 isoform in the viral life cycle and offer new mechanistic insights into EMCV translation.

## MATERIALS AND METHODS

### CRISPR screen experiment

The full-genome GeCKO v2 knockout pooled library, a gift from Feng Zhang (Addgene #1000000048; (71)), was packaged into lentiviral particles and used to transduce wild-type (WT) HEK293T cells at a multiplicity of infection (MOI) of 0.3. Selection of the transduced cells was performed using 0.5 μg/mL puromycin (twice the minimum lethal concentration) for 5 days. Subsequently, the selected cells were infected with EMCV at an MOI of 2. Following infection, we observed nearly complete cell death by the next day; however, two weeks later, single colonies of surviving cells began to emerge. Genomic DNA was isolated from these colonies and used as a template for the two rounds of PCR according to the protocol described earlier (72). The PCR products were sequenced on an Illumina NextSeq platform, and the resulting reads were analyzed using MAGeCK software (73).

### Cell culture and *ITAF45* knockout cell line production

HEK293T cells were cultured at 37 °C in a humidified atmosphere containing 5% CO_2_ in DMEM medium with 4.5 g/L glucose (PanEco, Russia), supplemented with 10% FBS (Capricorn Scientific, Germany), 1x GlutaMax (Gibco, USA), and 1x Penicillin-streptomycin antibiotic mixture (Gibco, USA). Once the cell density reached confluency, the cells were detached using 0.05% trypsin-EDTA (PanEco, Russia). To generate an *ITAF45* knockout (KO), we cloned a pair of oligonucleotides (5’-CACCGCTCCCCCCTTTTTGAAGAGGACCGACC-3’ and 5’-AAACGGTCGCTCGCTCTCTTCAAAAGGGGAGC-3’) into the px458 vector (Addgene plasmid #48138, a gift from Feng Zhang) via the BbsI restriction site. HEK293T WT cells were transfected with the knockout construct using PEI MAX (Polysciences, USA), employing a ratio of 3 μg of plasmid DNA to 9 μl of PEI (10 mg/ml) for each well of a 6-well plate. On the third day post-transfection, GFP-expressing cells were selected from the transfected population using a cell sorter (FACSAria SORP, BD Biosciences, provided by the Moscow State University Development Program) and subsequently cultured in a 6-well plate until reaching 80% confluency. The cells were then seeded onto a 96-well plate at a density of approximately 0.8 cells per well to establish monoclonal cell lines.

Western blotting was used to identify *ITAF45* KO cell lines (ITAF45-KO). Cultured cells were washed with phosphate-buffered saline (PBS), and proteins were extracted using lysis buffer (50 mM Tris-HCl pH 8.0, 150 mM NaCl, 0.1% SDS). SDS-PAGE was performed according to the manufacturer’s guidelines (Bio-Rad Laboratories, USA). Proteins were transferred to PVDF membranes using the Trans-Blot Turbo Transfer System and PVDF Transfer Packs (Bio-Rad Laboratories, USA). The membranes were then hybridized with a 1:1000 dilution of primary antibody against ITAF45 (66055-1-Ig, Proteintech Group, USA) and a 1:5000 dilution of goat anti-mouse secondary antibody (Invitrogen, USA) using the iBind Flex Western Device (Invitrogen, USA). Visualization was performed using Western ECL Substrate and the ChemiDoc imaging system (Bio-Rad Laboratories, USA). To analyze mutations at the target locus, PCR was performed using genotyping primers (5’-TGTAATGACGCAGTCCCCACC-3’ and 5’-TGGCAGTGCA-TTCCCTCTACC-3’) on genomic DNA extracted from the cells. The PCR products were separated by electrophoresis in agarose gel and sequenced by Sanger sequencing (Evrogen, Russia).

### Cell lines expressing p42 and p48 isoforms

Cell lines constitutively expressing p42 and p48 protein isoforms were generated based on the ITAF45-KO clone. For this purpose, we amplified DNA fragments encoding the p42 and p48 isoforms using cDNA synthesized from total RNA extracted from HEK293T cells. For each PCR reaction, we used a common primer for both isoforms, PA2G4-p42-p48-R: 5’-CACTCGGATCCGGATCCGTCCCCAG-CTTCATTCATTTTTTTTCT-3’, and specific primers for each isoform: PA2G4-p42-F: 5’-GCACTGGGGGATCCCCGCCA-CCACCATGATTATGGATGGAAGAAA-3’ or PA2G4-p48-F: 5’-ACGTGTGAGGATCCGCCGCCACCATGATGTCGGGCG-AGG-3’.

The resulting PCR products contained BamHI restriction sites at their 5’ and 3’ ends, allowing cloning into the lentiviral vector pHR-SFFV-KRAB-dCas9-P2A-mCherry (a gift from Jonathan Weissman, Addgene plasmid #60954), replacing the *dCas9* gene with the genes encoding the ITAF45 isoforms. The CDS of each isoform was followed by the coding sequence of the P2A peptide and the mCherry protein, enabling visualization of transgene expression in the cells. Thus, the transgenic proteins p42 and p48 carry an additional P2A peptide sequence at their C-terminus.

For lentiviral production, HEK293T cells were cotransfected with the ITAF45-encoding lentiviral vectors along with the psPAX2 and pMD2.G plasmids (a gift from Didier Trono, Addgene plasmids #12259 and #12260) using PEI (Polysciences, USA). Supernatants containing lentiviral particles were collected 48 h post-transfection (hpt) and filtered through a 0.20 μm filter (Merck, USA). ITAF45-KO cells were transduced with the lentiviruses at an MOI of 0.2. Four days after infection, mCherry-positive cells were selected using a cell sorter (FACSAria SORP). The sorted cells were cultured in a well of a 6-well plate until reaching approximately 80% confluency, then seeded into wells of a 96-well plate to establish monoclonal cell lines. Candidate clones, ITAF45-KO+p42 and ITAF45-KO+p48, were analyzed for homogeneity of the mCherry marker expression using the SinoCyte X flow cytometer (BioSino, China, provided by the Moscow State University Development Program). The expression levels of transgenic proteins in these candidate cell lines was assessed by Western blot, as described above.

### Mengo virus infection

We used the Mengo strain of EMCV from the collection of the FSBSI “Chumakov FSC R&D IBP RAS” to perform infection experiments. Virus was produced by infecting 1x10^7^ HEK293T cells at approximately 70% confluence. The following day, the viral-containing supernatant was filtered through a 0.20 μm filter (Millipore, USA). The resulting virus preparation was then frozen and stored at −20 °C. In all experiments, cells were infected with virus at an MOI of 2. Two hours post-infection, the medium was replaced with fresh medium to remove free virus not bound to the cells. During the course of infection, samples of the virus-containing medium were collected and stored at −20 °C. Viral titers were determined by infecting HEK293T cells plated in 96-well plates with serial dilutions of the virus-containing medium. The day after infection, cytopathic effect was visually assessed in the wells.

For the resazurin assay, cells were seeded in the wells of a 6-well plate at a density of 1×10^6^ cells per well. Prior to infection, the medium was replaced with 2 mL of fresh medium. The cells were infected with virus at an MOI of 2. After 24 h, 200 μL of a 440 μM resazurin solution was added to the cells. The cells were incubated in a CO_2_ incubator for 4 h, after which the wells were photographed.

The amount of EMCV RNA was measured by RT-qPCR. For this purpose, the culture medium was removed, and infected cells were lysed directly in the well by adding RNA Extract Solution (Evrogen, Russia), following the manufacturer’s protocol for RNA isolation. We used 2 μg of total RNA and an oligo(dT) primer to synthesize cDNA in a reverse transcription reaction with MMLV revertase (Evrogen, Russia). qPCR was performed using 5x qPCRmix-HS SYBR reagent (Evrogen, Russia) and the CFX96 Touch Real-Time PCR Detection System (Bio-Rad Laboratories, USA). Primers targeting the Mengo IRES region were: EMCV-qPCR-F1: 5’-TGAATGTCGTGAAGGAAGCAG-3’ and EMCV-qPCR-R1: 5’-CCCCTTGTTGAATACGCTTG-3’. Primers for the human *ACTB* gene (encoding β-actin) were: actin-F: 5’-GGCATCCACGAAACTACCTT-3’ and actin-R 5’-AACAGTCCGCCTAGAAGCAT-3’. PCR amplification conditions were as follows: initial denaturation at 95 **°**C for 2 min, followed by 40 cycles of 95 **°**C for 15 s, an annealing step 60 **°**C for 15 s, and extension at 72 **°**C for 15 s. Results were analyzed in the CFX Manager software (Bio-Rad Laboratories, USA). Each PCR reaction was performed in four technical replicates.

### Experiments with *in vitro* transcribed Mengo genomic RNA and Mengo replicon

The full-length Mengo genomic cDNA was cloned via RT-PCR using total RNA isolated from infected HK293T cells. For this, total cDNA was first synthesized with oligo(dT) primer and Magnus revertase (Evrogen, Russia). Subsequently, two fragments of the Mengo genome were amplified with specific primers: EMCV-CL-F: 5’-TTGAAAGCCGGGGGTGGGAGATCC-3’ and EMCV-3R: 5’-GTCATCCCTGCAATCAGCTGCACACATCT-3’ (fragment A); and EMCV-1F: 5’-TACGGCTTTGCTCGATGCCAACGAGGAC-3’ and EMCV-END-R: 5’-TTTTTTTTTTTTTTYWWCTATTT-ATTTTACTACTCTAGWC-3’ (fragment B). These fragments were cloned separately into the SmaI site of the pUC19 vector. Finally, Fragment A was excised and ligated into fragment B-containing plasmid using BstAFI and HindIII restriction sites, resulting in the pMengoWT plasmid.

Next, a plasmid encoding the Mengo replicon (pMengoFluc) was constructed in two steps. First, the *VP3-VP1* region was replaced with SpeI and NotI restriction sites by amplifying the entire pMengoWT plasmid with primers Spe-Not-VP1: 5’-CCACAACAC-TAGTGGGCGCGGCGGCCGCGCAGGAGTTCTTATGCTTGAA AGCC-3’ and Spe-VP2: 5’-CCAGCGACTAGTGATGGTCACA-GGGATAGGAGACTGTC-3’, followed by self-ligation. Then, the firefly luciferase (*Fluc*) coding region was amplified from the pGL3 vector (Promega, USA) with primers Spe-FLUC-F: 5’-CCAACAACTAGTGAAGACGCCA-AAAACATAAAGAAAGGC-3’ and Not-FLUC-R: 5’-CCACA-CGCGGCCGCACACGGCGATCTTTCCGCCCTTCTTG-3’, and inserted into the plasmid digested with SpeI and NotI.

For *in vitro* transcription, templates were generated from pMengoWT and pMengoFluc plasmids via PCR using the primers T7-CL-F-EMCV: 5’-AATTCTAATACGACTCACTATAGGGTTGAAAGCCGGGGGTGGGAGATCC-3’ and EMCV-ENDR-40A: 5’-(T)_40_CTATTTATT-TTACTACTCTAGT-3’. The uncapped transcripts were synthesized using the HiScribe T7 High Yield RNA Synthesis Kit (New England Biolabs, USA), precipitated using 2M LiCl, and checked for integrity via denaturing urea-polyacrylamide gel electrophoresis, as described previously (74). The transcripts were used to transfect HEK293T cells according to a protocol described below. In the case of MengoWT RNA, gRNA abundance was quantified by RT-qPCR at various time points post-transfection. For the replicon, Fluc activity was measured in cell lysates, as detailed in the mRNA transfection section.

### Recombinant ITAF45 protein purification

The CDSs of the p42 and p48 isoforms were cloned into the pET28b(+) vector using BamHI and XhoI restriction sites. The previously described lentiviral constructs encoding the isoforms as templates, along with the following primers, were used for PCR amplification: p42-F: 5’-CCAACTGGATCCTATTATTATTATGGATGGAAGAAAC-AGGGAAACAGGGAAAATCTTCAAGAAAG-3’, p48-F: 5’-GCAACACAGGATCCTTCGGGCGAGGACGACGACGAGCAAC AGG-3’, p42-48-R: 5’-CCAATACTCGAGTTAGTTAGTAGTCC-CCAGCTTCTTCATTCATTTTTTCTTCTTCTTCTAATGTTTTTCC-3’. *E. coli* Rosetta strain was used for protein production. A single colony of plasmid-transformed cells was transferred into a small volume of LB medium supplemented with kanamycin and chloramphenicol, and grown with constant shaking at 37 **°**С. The culture was then diluted into a larger volume (500 mL) of medium and grown until reaching an OD600 of 0.6-0.7. The culture flasks were cooled down to 18 °C, and expression was induced by adding IPTG to a final concentration of 0.5 mM. Cultures were incubated overnight at 18 °C with constant shaking. Subsequently, cultures were centrifuged at 5,000 g at +4 °C, the medium was removed, and the bacterial pellet was resuspended in PBS-based buffer supplemented with 10 mM imidazole, 0.5 mM PMSF protease inhibitor, and 10% glycerol. Cells were lysed by ultrasonication. The lysate was centrifuged at 20,000 g at +4 °C, then 3M KCl was slowly added to the supernatant to reach a final concentration of 300 mM. The mixture was combined with Ni-Azur resin equilibrated with the same buffer and incubated at rt with shaking for 30 min. The suspension was transferred to a gravity flow column, the flow-through was discarded, and the resin was washed with PBS-based buffer supplemented with 20 mM imidazole, 10% glycerol, 0.5 mM PMSF, and 300 mM KCl. Target proteins were eluted by adding portions of the same buffer containing 250 mM imidazole, the collected fractions were analyzed by SDS-PAGE and Coomassie R-250 staining. Fractions containing the target protein were pooled, diluted with buffer A0 (20 mM Tris-HCl pH 7.5, 3 mM β-mercaptoethanol, 10% glycerol, 0 mM KCl) to reduce the final KCl concentration to 100 mM, and further purified via FPLC ion-exchange chromatography on a monoQ column equilibrated with buffer A100 (20 mM Tris-HCl pH 7.5, 3 mM β-mercaptoethanol, 10% glycerol, 100 mM KCl). Proteins were eluted with a gradient increasing KCl concentration from 100 mM to 730 mM. Fractions were analyzed by SDS-PAGE followed by Coomassie R-250 staining. Fractions containing pure target protein were pooled and concentrated using Amicon Ultra 10 kDa centrifugal filters (Merck, USA), with simultaneous buffer exchange to A100.

### mRNA transfection of cultured cells, dual-luciferase assay, real-time luciferase measurement, and fluorescent flow cytometry

The plasmid encoding the bicistronic Rluc-EMCV_IRES-Fluc mRNA has been described previously (75). Similar plasmids containing the FMDV IRES and the full-length Mengo 5’ UTR sequences were generated as follows: FMDV C-S8 regions containing IRES with either AUG1 or AUG1 along with the beginning of the L protease coding sequence and AUG2 were amplified from pFMDVluc (24) using the primers FMDV-F: 5’-GCAGGTTTCCCCAACTG-ACAC-3’ and FMDV-AUG-lab-R: 5’-CTGACTTCCATGGCA-GGGTCATTAATTGTAAAGGAAAGGTG-3’, or FMDV-F: 5’-GCAGGTTTCCCCAACTGACAC-3’ and FMDV-AUG-lb-R: 5’-CGACTTCCATGGTTTTCCTG-CAGTCCGTGGTAG-3’. The amplified fragments were phosphorylated with PNK, digested with NcoI, and cloned into the NcoI and PvuII sites of the pGL3R vector (76), resulting in Rluc-FMDV_IRES-AUG1-Fluc and Rluc-FMDV_IRES-AUG2-Fluc. To obtain Rluc-MENGO_5’UTR-Fluc, the Mengo 5’ UTR was amplified from pMengoWT using primers Nco-EMCV-CL: 5’-CCACACCATGGTTGAAAGCCGGGGGTGGGA-GATCC-3’ and Nco-EMCV-5UTR-R: 5’-AAATCTCTTGTTC-CATGGTTGTAGCCAT-3’, digested with NcoI, and cloned into pGL3R. The fluorescent reporter KAT-EMCV-EGFP was created by inserting a PCR product, obtained from pTurboFP635-C (Evrogen) with the primers KAT-F: 5’-GCGACGGATCCGAAATAAGAGAGAAAAGAAGAGTAAGAAG-AAATATAAGAGCCAAGATGGTGGGTGAGGATAGCGTGC-3’ and KAT-R: 5’-GGCTCGAATTCTTTAGCTGTGCCC-CAGTTTGCTAGGCAGGTCGCAGTATCTAGCCACAGC-3’, into the BamHI and EcoRI sites of the LeGO-iG2 lentiviral vector (Addgene #27341, a gift from Boris Fehse). The construct was kindly provided by Alexander Konopatov (MSU). In line with MINDR recommendations (77), sequences of all plasmid are provided in Supplementary Data.

For *in vitro* transcription, T7 promoter-containing, 50A-tailed templates were generated via PCR from the respective plasmids, as described previously (78), using primers: RV3long: 5’-CTAGCAAAATAGGCTGTCCCCAG-3’ and FLA50: 5’-(T)_50_AACTTGTTTATTGCAGCTTATAATGG-3’ for the pGL3R-based plasmids, or with T7-R-region-F: 5’-CCTTCTAATACGACTCACTATAGGGCCAGTCCTCCGATTGAC TGAG-3’ and PolyT-LTR-PAS-R: 5’-(T)_50_GAAGCACTCAAG-GCAAGCTTTATTG-3’ for KAT-EMCV-EGFP. Reporter mRNAs were prepared as described above and co-transcriptionally capped with the ARCA cap analog (BioLabMix, Russia; 5:1 ARCA to GTP).

Cell transfection with luciferase mRNAs was performed according to the Fleeting mRNA Transfection (FLERT) protocol (74), which allows rapid and minimally stressful transfection. Briefly, cells were seeded into 24-well plates and cultured overnight to approximately 70% confluence. Transfection was carried out in duplicate using the GenJector-U reagent (Molecta, Russia). For luciferase assays, cells were harvested 6 hpt, and luciferase activities were measured using the Dual-Luciferase Reporter Assay System (DLR, Promega, USA) on a TD-20/20 Compact Luminometer (Turner Designs Instruments). All experiments were performed at least three times, including different cell passages, and the mean ± standard deviation (SD) values were calculated.

For flow cytometry analysis, cells were transfected with reporter RNA using roughly the same protocol, except that 6-well plates and 1 μg of mRNAs were used. Two days post-transfection, cells were detached with 0.05% trypsin-EDTA solution, and trypsin was neutralized with an equal volume of culture medium. The cell suspension was centrifuged for 5 min at 500 g. The cell pellet was resuspended in 200 µL of culture medium. Fluorescence was measured using a SinoCyte X flow cytometer (BioSino, China): eGFP fluorescence was detected in the FITC channel (525/40 nm), and TurboFP635 fluorescence was measured in the mPlum channel (660/20 nm). Transfections were performed in four replicates. Flow cytometry data were analyzed using FlowJo software, and results were visualized with the Floreada online service and GraphPad Prism.

An mRNA transfection followed by real-time measurement of Fluc activity was performed essentially as previously described (79). Briefly, one day before transfection, approximately 3×10^4^ cells per well were transferred into white FB/HB 96-well plates (Greiner, Austria) in 80 μl of DMEM supplemented with 10% FBS and penicillin-streptomycin mixture. The next day, when the cell culture reached approximately 70% confluency, transfection was performed with reporter mRNAs. For this, 30 ng of mRNA in 17.6 μl of Opti-MEM (Gibco, USA) per well was mixed with 0.04 μl of GenJector-U in 2 μl of Opti-MEM (per well), incubated for 15 min, then supplemented with 0.4 μl of 100 mM D-luciferin (Promega, USA) per well. Subsequently, 20 μl of the mixture were added to the cells. During these steps, all manipulations were performed as described in the FLERT protocol (74). Real-time luminescence measurements were carried out overnight in the CLARIOstar plate reader (BMG Labtech, Germany) equipped with an Atmospheric Control Unit to maintain 5% CO_2_ at 37 °C (signal integration time: 5 s). Transfection of both WT and KO cells with capped and polyadenylated Actin-Fluc mRNA (75) was performed in parallel in the same experiment. The resulting average values were used to normalize the data obtained from the wells with IRES-containing reporters, accounting for possible differences in cell density and transfection efficiency between the two cell lines. All experiments were repeated at least three times (including those with different cell passages), and mean values ± SD were calculated.

### Translation in cell-free systems

For *in vitro* translation, whole-cell extracts prepared from cultured cells of the indicated cell lines were used as described earlier (17), with some modifications (80). Briefly, cells were grown in 150-mm culture dishes to approximately 75% confluency; 30 min before harvesting, 1/200х of 100x MEM Non-Essential Amino Acids Solution (Gibco, USA) was added to the medium. Cells were then incubated on ice for 5 min, the medium was aspirated, and cells were washed twice with ice-cold DPBS+G (Dulbecco PBS containing 25 mM D-glucose), followed by scraping the cells with a cell lifter (3008, Corning, USA) into 7 mL of fresh DPBS+G. Cells were centrifuged for 5 min at 600 g, resuspended in 3 mL of DPBS+G, transferred to 1.5 mL tubes, and centrifuged for 5 min at 600 g. The pellet was then resuspended in Hypotonic Extraction Buffer (25 mM Hepes-KOH pH 7.6, 0.5 mM Mg(OAc)_2_, 10 mM KOAc, 2 mM DTT) in a 1:1 ratio to the cell volume and incubated on ice for 30 min. Finally, the cells were lysed by 20-25 gentle strokes with a 2 mL insulin syringe fitted with a 26G needle and centrifuged for 10 min at 15,000 g at 4 °C. The middle fraction of the supernatant was collected, aliquoted, flash-frozen in liquid N_2_, and stored at -85 °C.

Translation reactions were performed in a total volume of 10 μL, containing 5 μL of the cell extract, 1x Translation Buffer (20 mM Hepes-KOH pH 7.6, 70 mM KOAc, 1 mM Mg(OAc)_2_, 1 mM DTT, 0.5 mM spermidine-HCl, 8 mM creatine phosphate, 1 mM ATP, 0.2 mM GTP, and 30 μM of each amino acid), 2 u of RiboLock RNase inhibitor (Thermo Fisher Scientific, USA), 1 μL of either A100 buffer or protein solution (as indicated), and 15 ng of mRNA (added as 1 μL water solution after the pre-incubation of the reaction mixture for 5 min at 30 °C). After mRNA addition, the mixtures were transferred to 0.5-mL tubes, incubated for 30 min at 30 °C, and placed on ice. Fluc and Rluc activities were measured from 2.5-μL aliquots of the reactions by sequential addition of 10 μL LAR II and 10 μL Stop & Glo buffers from the DLR kit (Promega, USA) on a TD-20/20 Compact Luminometer. All experiments were performed in duplicate and repeated at least twice; results are presented as mean ± SD.

### Statistical analysis

Statistical analyses were performed using GraphPad PRISM 8.0.1 software. A p-value of less than 0.05 was considered statistically significant. Data from two groups were analyzed using an unpaired t-test, while data involving more than two groups were assessed using either one-way or two-way ANOVA, followed by Tukey’s post hoc test to identify specific group differences. The line-of-best fit was determined using GraphPad Prism, and the slopes of the lines were compared through linear regression analysis. Data are presented as the mean ± SD of at least three independent experiments.

## RESULTS

### ITAF45 is an essential cellular factor for cytopathic EMCV/Mengo infection

To identify host genes involved in the progression of EMCV/Mengo infection, we conducted a genome-wide knockout CRISPR screen in human embryonic kidney HEK293T cells, which exhibit cytopathic effects upon viral infection. Among the genes whose knockout potentially promotes cell survival following virus infection (Supplementary Table S1), we found *ADAM9*, which has previously been implicated in EMCV entry into the cell (81,82). Intriguingly, we also found the *PA2G4*/*EBP1ITAF45* gene at the top of the list; its product, hereafter called ITAF45, has been shown to be essential for the assembly of the 48S pre-initiation complex on the FMDV IRES but was reported to have no effect on translation initiation directed by the EMCV IRES (28,62). Further analysis of data from other CRISPR screen experiments conducted with EMCV, available in the BioGRID ORCS database (83), revealed that *PA2G4* consistently ranked among the top hits across nearly all studies. It was also identified as the top hit in two of the most comprehensive CRISPR screens of EMCV infection (81,82), although this observation has not been further investigated. This raises important questions regarding the necessity of ITAF45 in the EMCV/Mengo lifecycle and its role in IRES-dependent translation.

To validate the hypothesis that ITAF45 is required for the progression of EMCV/Mengo infection, we generated a monoclonal cell line with a CRISPR/Cas-induced knockout of the *PA2G4*/*ITAF45* gene, derived from HEK293T cells. Western blot analysis confirmed the complete absence of the ITAF45 protein in the knockout (KO) cells (Figure 1A). Genotyping of the KO clone revealed nucleotide deletions at the gRNA-targeted genomic locus: an 8-nt deletion in one allele and a 131-nt deletion in the other, which disrupts the junction between the second intron and the third exon of the gene (Figure 1B, Supplementary Figure S1).

**Figure 1.**
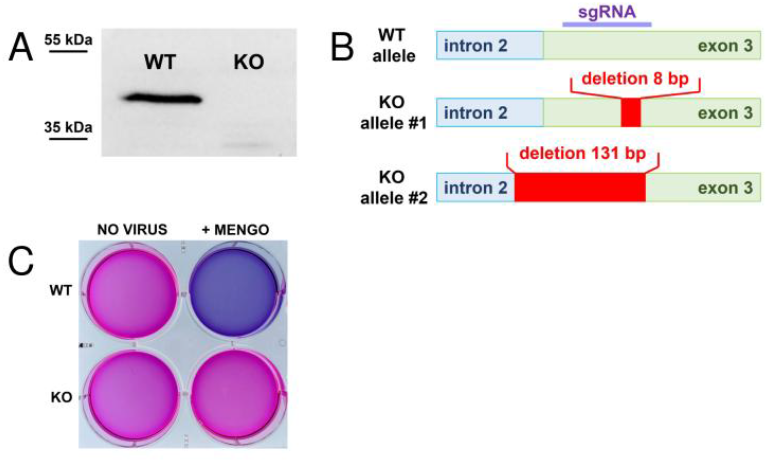
ITAF45 is essential for EMCV/Mengo virus infection in human cells. (А) Verification of ITAF45 protein absence in the *ITAF45* KO HEK293T clone by Western blot. (B) Schematic representation of mutations in the alleles of the *ITAF45* gene in the KO clone. The sgRNA was targeted to the beginning of the third exon. In the first allele, an 8 bp deletion occurs at the start of exon 3, which causes a frameshift. In the second allele, an extended 131 bp deletion spans the exon/intron junction, disrupting the splicing site region. (С) HEK293T WT and ITAF45-KO cells treated with resazurin solution. The top row shows uninfected cells, while the bottom row shows cells 24 h after Mengo infection (MOI = 2). WT cells were completely destroyed by the virus, whereas KO cells survived the infection.

The KO cells survived Mengo infection (MOI = 2) at 8 h post-infection (hpi) and remained fully viable 24 hpi, whereas wild-type cells exhibited a pronounced cytopathic effect at 8 hpi and were completely dead after 24 h (Figure 1C). Thus, our findings indicate that ITAF45 is required for cytopathic Mengo infection in HEK293T cells.

### ITAF45 p48 but not p42 isoform supports productive EMCV/Mengo infection

In human cells, the ITAF45 protein exists in two isoforms, p42 and p48, which arise from alternative pre-mRNA splicing (Figure 2A). These isoforms exhibit differential expression patterns in normal and cancerous tissues (as noted previously). The p42 isoform is characterized by the absence of the N-terminal α-helix and a portion of the second α-helix (Figure 2B), which significantly impacts its functional capacity and likely contributes to its instability. Since both isoforms localize to the cell cytoplasm, where the Mengo life cycle takes place, we aimed to evaluate the ability of each isoform to maintain viral infection.

**Figure 2.**
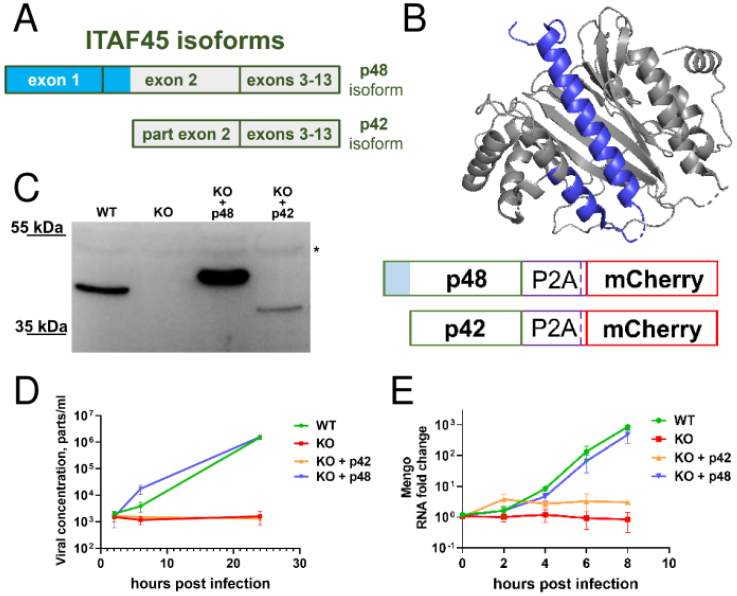
Differential activity of ITAF45 isoforms in supporting productive EMCV/Mengo virus infection in human cells. (A) A schematic of the two ITAF45 isoforms. The longer p48 isoform is synthesized from an mRNA containing 13 exons, whereas the shorter p42 isoform originates from an mRNA lacking the first exon. (B) Three-dimensional model of ITAF45 (PDB ID 2Q8K; (61)). The N-terminal α-helix 1 and a portion of α-helix 2 (both colored blue) are present only in the p48 isoform. (C) Western blot analysis of ITAF45 protein levels in KO cells expressing either the p42 or p48 isoform. The reduced mobility of the transgenic proteins can be explained by the presence of a P2A peptide sequence at the C-terminus of the protein (right panel). ITAF45-KO+p42 cells show lower protein levels, likely due to intrinsic instability. A nonspecific band is marked with an asterisk. (D) Dynamics of virus production in infected cells. WT, KO, and KO cells exclusively expressing the p42 or p48 isoforms were infected with Mengo virus (MOI = 2), and at the indicated times after infection the concentration of viral particles in the culture medium was determined by a serial-dilution method. Note the log scale of the Y axis. Experiments were performed in three biological replicates. The means ± SD are shown. (E) Dynamics of Mengo gRNA accumulation in infected cells. WT, KO, and KO cells exclusively expressing the p42 and p48 isoforms were infected with Mengo virus (MOI = 2), and the accumulation of viral RNA was analyzed at different time points by RT-qPCR. Note the log scale of the Y axis. Experiments were performed in three biological replicates, with qPCR reactions performed in four technical replicates. The means ± SD are shown.

To investigate this, we transduced the ITAF45-KO cell line with lentiviral constructs that constitutively express either the p42 or p48 isoform and subsequently generated monoclonal cell lines. Western blot analysis confirmed the successful production of the transgenic p42 and p48 proteins, which exhibited slightly higher molecular weights than anticipated due to the inclusion of the C-terminal P2A peptide (Figure 2C). Notably, despite using the same expression cassette and similar levels of *mCherry* marker expression (Supplementary Figure S2), p42 expression was lower than p48, likely reflecting the intrinsic instability of the p42 isoform. However, p42 abundance was comparable to endogenous p48 in WT cells. Interestingly, WT HEK293T cells express only the p48 isoform.

We then assessed the ability of Mengo virus to propagate in WT, KO, and KO cells expressing either the p42 or p48 isoforms (Figure 2D). WT cells and cells selectively expressing the p48 isoform effectively produced infectious virus particles. In these cells, Mengo infection induced pronounced cytopathic effects, leading to complete cell death by 24 hpi (data not shown). In contrast, KO cells and those selectively expressing the p42 isoform showed no increase in the number of active virus particles or cytopathic effects.

To elucidate whether the observed effects were due to inability of the virus to replicate in KO and p42-expressing cells, we conducted RT-qPCR analysis to assess genomic RNA accumulation. The growth of viral RNA concentrations in cells expressing the p48 isoform mirrored that of WT cells, demonstrating similar dynamics. In contrast, p42-expressing cells either accumulated genomic RNA very inefficiently or failed to accumulate it at all, similarly to the KO cells (Figure 2E).

Collectively, these experiments indicate that the p48 isoform of ITAF45 is essential for Mengo infection in cultured cells, whereas the p42 isoform is insufficient for this purpose.

### ITAF45 KO cells transfected with synthetic Mengo genomic RNA or RNA replicon fail to support their replication

The entry of viruses into host cells is a complex, multistage process that requires the involvement of numerous cellular proteins. The observed dependence of Mengo virus on ITAF45 for successful infection may, in theory, be indirect, potentially mediated by cellular proteins whose production or activity requires the *ITAF45* gene product. To investigate this possibility, we cloned the full-length Mengo genome and generated artificial full-length genomic RNA through *in vitro* transcription. Additionally, we constructed a replicon by replacing the structural protein genes *VP3* and *VP1* with the *Fluc* gene, while retaining regions corresponding to several N- and C-terminal amino acid residues of the VP3 and VP1 to facilitate the efficient excision of the luciferase protein from the synthesized polypeptide by viral proteases, and synthesized the resulting transcript (Figure 3A).

**Figure 3.**
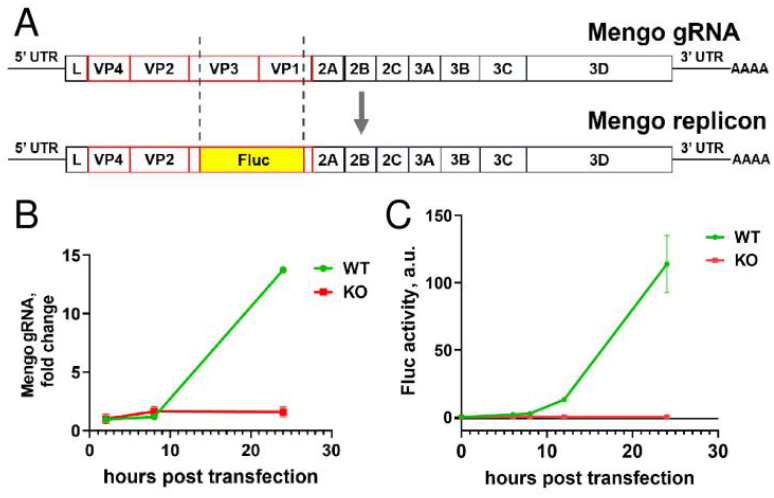
ITAF45-KO cells transfected with synthetic Mengo RNAs fail to support their replication. (A) A schematic representation of the Mengo genomic RNA (top) and Mengo replicon RNA in which the *VP3* and *VP1* capsid protein genes are replaced with the *Fluc* reporter gene (bottom). (B) Mengo gRNA accumulation in WT and KO cells transfected with *in vitro*-transcribed viral gRNA, measured at 2, 8, and 24 hpt by RT-qPCR. The experiments were conducted in three biological replicates, with qPCR reactions performed in four technical replicates. The means ± SD are shown. (C) Luciferase activity in WT and KO cells at 6, 8, 12, and 24 hpt with Mengo replicon RNA. The means ± SD of three technical replicates are shown.

The *in vitro* synthesized full-length genomic RNA delivered to WT cells via lipofection successfully replicated, as evidenced by a sharp increase in RT-qPCR product levels, escalating by two orders of magnitude at 24 hpt (Figure 3B). In stark contrast, the ITAF45-KO cells exhibited no accumulation of viral RNA. Similar results were obtained with the replicon lacking the VP3-VP1 region (Supplementary Figure S3), further supporting the hypothesis that ITAF45 dependence is not related to viral particle production or cell entry.

Next, we assessed luciferase expression in both WT and KO cells transfected with the replicon RNA. A notable tenfold increase in luciferase activity was detected in WT cells compared to KO cells as early as 6 hpi, with this difference amplifying to three orders of magnitude by 24 hpt (Figure 3C).

Collectively, these results clearly indicate that ITAF45 is essential at certain stage(s) following viral entry into the cell.

### The role of the p48 isoform of ITAF45 in the activities of EMCV/Mengo and FMDV IRESs

In an earlier study, ITAF45 was identified as essential for the assembly of the 48S pre-initiation complex from purified components on the FMDV IRES, while it was not required for assembly on the EMCV IRES (28). We hypothesized that ITAF45 could still play a role in the translation of EMCV/Mengo virus RNA, albeit at a different stage or under specific conditions.

To test this hypothesis, we generated bicistronic reporter constructs in which the translation of the first cistron (*Renilla luciferase, Rluc*) was cap-dependent, while the second cistron (*Fluc*) was under the control of either FMDV or EMCV IRESs, or the full-length Mengo 5’ UTR (Figure 4A). It should be noted that the “EMCV IRES” in this construct corresponds to that used in the original study by Pestova et al. (27). Then, the EMCV and Mengo nucleotide sequences were very similar in their IRES-containing region (∼540-nt AUG-proximal segment), while the full-length Mengo 5’ UTR harbored an additional 5’ proximal region including a poly(C)-tract. For FMDV, we prepared two constructs with the *Fluc* CDS starting from either AUG1 or AUG2 of the FMDV gRNA (see Introduction). However, in all our experiments these two constructs yielded very similar results, so hereafter the term “FMDV” denotes the Rluc-FMDV_IRES-AUG2-Fluc construct. Experiments utilizing these bicistronic constructs were performed in accordance with the minimal dual reporting requirements for experiments involving dual gene expression reporters (MINDR) (77).

**Figure 4.**
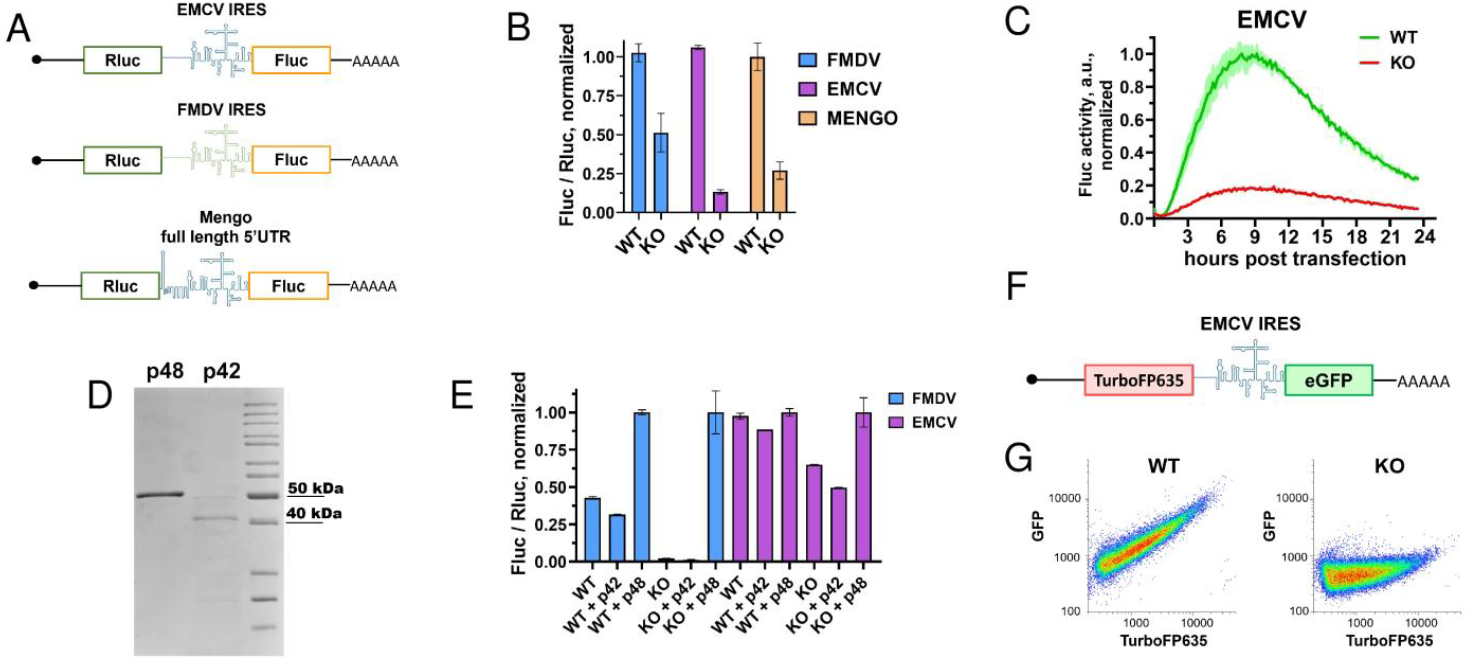
ITAF45 p48 is needed for EMCV and FMDV IRES activities in cultured cells and cell-free systems. (A) A schematic representation of the capped and polyadenylated bicistronic luciferase mRNAs used. (B) Normalized luciferase activities in WT and KO cells transfected with the bicistronic mRNAs shown in (A). Fluc/Rluc values reflect IRES activity. Data are presented as means ± SD. (C) Time-course curves of real-time Fluc activity in WT and KO cells transfected with the Rluc-EMCV_IRES-Fluc mRNA shown in (A). Fluc values were normalized to the average Fluc value in wells of the same cell line transfected with capped and polyadenylated Actin-Fluc mRNA (to account for any non-specific differences between the cell lines) and then to the maximal value in WT. SDs are shown as semi-transparent shading. (D) SDS-PAGE of the purified recombinant p42 and p48 proteins. (E) EMCV- and FMDV IRES-driven reporter activities in cell-free translation systems supplemented with 0.5 µg of each recombinant protein. Fluc and Rluc activity values were measured at 30 min, Fluc/Rluc ratios were calculated and normalized to those in “+p48” experimental condition with the corresponding constructs and lysates (to account for non-specific differences between WT and KO lysates). Experiments were performed in two biological and two technical replicates. The means ± SD are shown. (F) A schematic of the capped and polyadenylated bicistronic fluorescent reporter mRNA. (G) Flow cytometry data for WT and KO cells transfected with the KAT-EMCV_IRES-EGFP mRNA. The analysis was performed 24 hpt. Experiments were performed in three biological replicates; a representative example is shown in (G), while all results are available in Supplementary Figure S4.

WT and KO cells were transfected with capped and polyadenylated transcripts following the Fleeting mRNA transfection (FLERT) protocol (74). Luciferase activities were analyzed 6 hpt. As expected, consistent with previous findings (28,62), the FMDV IRES-dependent translation was diminished in the ITAF45-KO cells compared to WT HEK293T cells, exhibiting approximately a two-fold reduction (Figure 4B). Notably, EMCV IRES activity demonstrated an even greater dependence on ITAF45, resulting in an approximately 8-fold decrease in reporter translation in KO cells.

In a separate experiment, we also applied another mRNA transfection protocol with real-time monitoring of Fluc activity (79), which provides data on mRNA expression at multiple time points. Using this assay, we observed a similar dependence of EMCV IRES activity (Figure 4C), which was roughly consistent throughout the entire transfection period.

Subsequently, we examined the translation activity of bicistronic mRNAs containing the EMCV and FMDV IRESs in cell-free systems prepared from WT and KO cells, supplemented with recombinant ITAF45 proteins (Figure 4D-E). These self-made cell lysates closely recapitulate translation activity in living cells but enable the direct addition of exogenous components (17,75). In this analysis, the FMDV IRES showed a substantially greater dependence on ITAF45 (Figure 4E), whereas the EMCV IRES exhibited only a modest decrease in activity (approximately two-fold) in the absence of ITAF45. We then took advantage of cell-free systems to directly introduce purified proteins. Recombinant p42 and p48 isoforms were produced and purified from *E. coli* (Figure 4D), and subsequently added to the cell lysates to facilitate translation of the EMCV- and FMDV IRES-containing bicistronic mRNAs in these systems. Notably, the addition of recombinant p48 protein restored the ability of the KO cell lysate to mediate EMCV- and FMDV IRES-dependent translation, whereas p42 had no positive effect (Figure 4E).

Finally, to eliminate potential biases arising from the choice of reporter type, we also created an alternative bicistronic construct encoding fluorescent proteins, where synthesis of TurboFP635 (Katushka) was cap-dependent while eGFP production was driven by the EMCV IRES (Figure 4F). We observed a direct correlation between the levels of TurboFP635 and eGFP in WT cells, whereas nearly no GFP expression was detected in KO cells (Figure 4G, Supplementary Figure S4). This result emphasizes the essential function of ITAF45 in supporting EMCV IRES activity across different reporter systems.

In summary, our findings demonstrate that ITAF45 p48 isoform is essential for the efficient functioning of the EMCV IRES in human cells and cell-free systems.

## DISCUSSION

This study reveals a previously unrecognized role of the p48 isoform of ITAF45/EBP1/PA2G4 in mediating EMCV infection, primarily by facilitating IRES-dependent translation of viral RNAs. Using knockout cell models, we establish ITAF45 as an indispensable host factor for efficient cytopathic infection by EMCV/Mengo virus and demonstrate that the p48 isoform of ITAF45 is specifically required for the viral life cycle, whereas the p42 isoform is insufficient to support this process. The failure of viral replication upon transfection of synthetic Mengo genomic RNA or replicons into ITAF45-KO cells suggests that ITAF45 functions at a post-entry stage, likely during translation or replication initiation. Moreover, analyses of IRES activity revealed that the p48 isoform is vital for the optimal function of the EMCV IRES in cultured cell.

It was previously reported that ITAF45 is essential for the assembly of the 48S pre-initiation complex from purified components on the FMDV IRES (28) but dispensable in the case of two related IRESs, TMEV (Theiler’s encephalomyelitis virus) and EMCV (27,28). It was also claimed that addition of recombinant mouse ITAF45/Mpp1 (ortholog of the human ITAF45 p48 isoform) did not influence the 48S complex formation on the EMCV IRES (stated in (28) as data now shown). In partial accordance with that, in the complete translation system prepared from cultured human cells, we found strong dependence of the FMDV IRES-driven translation (initiated from both AUG1 and AUG2), whereas much less prominent dependence of the EMCV IRES activity.

In contrast, in living cells, our data indicate that efficient expression of EMCV/Mengo RNA clearly requires ITAF45. This contradicts the earlier study by Monie et al. (62). The discrepancy could reflect differences in experimental design: although the other researchers also transfected human HEK293 cells with bicistronic mRNAs, they used RNA interference instead of gene knockout and employed a different mRNA transfection protocol. Specifically, they sequentially transfected cells with siRNA (for 48 h) and then with mRNA reporters (for an additional 24 h). Based on our experience (see e.g. (79,84)), a 24-h incubation period after mRNA transfection may be too long, as illustrated by the curves in Figure 4C. Moreover, performing two sequential transfections could be toxic to cells and potentially influence the results. Furthermore, in our study, the requirement of EMCV for *ITAF45* in living cells is confirmed through multiple approaches: it is observed both with viral gRNA and replicon, as well as in the context of various mRNA reporters, across multiple time points. Our findings are also consistent with direct RNA-protein interaction assays showing that ITAF45 binds to the EMCV IRES (28,30,62).

The fact that the dependence of activities of both EMCV/Mengo and FMDV IRESs varies among different systems raises questions about the unambiguous role of ITAFs in IRES function. Such varying requirements for ITAFs, depending on the cellular context (host cell type, stress conditions, or experimental system), as described in earlier studies (reviewed in (11,85)), underscore the complexity of IRES-mediated translation regulation and the importance of context-specific ITAF interactions. They also highlight the adaptability of IRESs to different host environments (e.g., neurovirulence of the virus) and their complex interplay with cellular regulatory pathways. Moreover, the observed discrepancy between EMCV IRES dependence on ITAF45 in living cells and in translation lysates prepared from the same cell lines suggests that ITAF45 may participate in processes beyond initiation complex formation, such as regulation of elongation, termination, or ribosome recycling; localized translation; or mRNA quality control. Further investigation is needed to clarify these issues.

The findings from our study underscore the specific role of the p48 isoform of ITAF45 in facilitating EMCV and FMDV IRES activity, while the N-terminally truncated p42 isoform was found to be inactive. It is important to note that, in complex with the mammalian ribosome, ITAF45 binds simultaneously to several rRNA segments (59,63,65). Such simultaneous binding to multiple RNA domains is likely a general feature of ITAFs, which act as RNA chaperones to stabilize IRESs in their active conformation (30-32). Among the rRNA segments interacting with ITAF45 in its ribosomal complex, the long double-stranded helix of ES27L is of particular interest. ITAF45 interacts with ES27L through two distinct regions: (i) the two N-terminal α-helices, and (ii) the C-terminal Lys-rich region, which has also been shown to be important for the interaction of mouse ITAF45 with the FMDV IRES (62). The inability of p42, which lacks the first α-helix and part of the second α-helix, suggests that the RNA-binding surface formed by these two regions is critical for the protein’s ITAF function.

When this manuscript was in preparation, a preprint by Bellucci et al. was published (86), demonstrating that ITAF45 is an essential host factor for four picornaviruses containing Type II IRESs, including EMCV, and that it enhances IRES activity in cultured cells. They also showed that the C-terminal Lys-rich region (residues 365–376) is indispensable for this activity.

Collectively, these two findings position the p48 isoform of ITAF45 as a crucial modulator of picornaviral IRES-driven translation, which may have significant implications for understanding the translational strategies employed by viral pathogens. They also emphasize the critical importance of the RNA-binding patch, comprising two N-terminal α-helices and the C-terminal Lys-rich region, for this function. Future studies focusing on the mechanistic details of how ITAF45 interacts with viral IRES elements will provide a more comprehensive understanding of its ITAF activity, the regulation of viral gene expression, and its impact on pathogenesis. Given the essential role of ITAF45 in IRES-mediated translation of picornaviral RNAs, further investigation into its role during viral infection may reveal novel therapeutic targets for antiviral strategies.

## Supporting information

Supplementary Data - Plasmid Maps

## CONFLICT OF INTEREST

The authors have no conflicts of interest to disclose.

## SUPPLEMENTARY DATA

Supplementary Data are available at BioRxiv.

## AUTHOR CONTRIBUTIONS

Conceptualization, V.I.A. and S.E.D.; data curation, A.S.K., V.A.G., E.A.P., A.P.S., and S.E.D.; formal analysis, A.S.K., V.A.G., E.A.P., A.P.S., and S.E.D.; funding acquisition, S.E.D.; investigation, A.S.K., V.A.G., E.A.P., A.P.S., E.E.G., Y.Y.I., A.Y.K., and S.E.D.; methodology, A.S.K., V.A.G., E.A.P., A.P.S., A.Y.K., A.V.P., and S.E.D.; project administration, S.E.D.; resources, A.V.P., Y.Y.I., and S.E.D.; software, A.S.K., V.A.G., E.A.P., and A.P.S.; supervision, V.I.A. and S.E.D.; validation, A.S.K., V.A.G., E.A.P., A.P.S., and S.E.D.; visualization, A.S.K. and V.A.G.; writing – original draft, A.S.K. and S.E.D.; writing – review and editing, A.P.S., E.A.P., and S.E.D. All authors have read and agreed to the published version of the manuscript.

## FUNDING

This work was conducted under the state assignment of Lomonosov Moscow State University and was supported by the Russian Science Foundation (grant no. 23-14-00218 to S.E.D.)

## ACKNOWLEDGMENTS

We are grateful to Artem Ferberg (MSU, Russia), Olga Khudoleeva (Helicon, Russia), and Daria Potashnikova (MSU) for help with flow cytometry experiments; Margarita Ezhova and Maria Logacheva (MSU) for CRISPR-screen amplicon sequencing on the Illumina platform; Alexander Konopatov (MSU and IGB RAS, Russia) for the KAT-EMCV-EGFP plasmid; and Ivan Sorokin (MSU, Russia; University of Groningen, the Netherlands) for valuable suggestions regarding cell-free systems. We are also grateful to the Moscow State University Development Program for providing access to the FACSAria SORP cell sorter and the SinoCyte X flow cytometer.

## Notes

### Competing Interest Statement

The authors have declared no competing interest.

### Summary of Updates

We added additional Supplementary Figure and fixed some minor textual bugs

