## Supplementary Table and Figures for "The p48 isoform of the PA2G4/EBP1/ITAF45 oncoprotein is required for the encephalomyocarditis virus IRES-driven translation initiation"

### SUPPLEMENTARY DATA

---

**Supplementary Table 1.** Top six sgRNA hits from a genome-wide knockout CRISPR screen in HEK293T cells identifying host genes involved in EMCV/Mengo virus infection. Read counts from the Illumina sequencing library are shown. *PA2G4* and *ADAM9*, as mentioned in the main text, are highlighted.

| Gene | Count | Sequence |
|---|---|---|
| *PA2G4* | 903 | CTCCCCTTTGAAGAGCGACC |
| *ADAM9* | 681 | TTAGGTATCTTATGTTATTC |
| *CSE1L* | 622 | CGTTTGACTTAAATTCATGA |
| *PAX7* | 320 | GCCGGATGGACCCGGTCTCC |
| *RREB1* | 239 | GACAATCGCCTACGTTTCAG |
| *CT47A9* | 93 | GCTGGTGTCATGTCTGCCAC |

```
WT        CTACATTTCTGAAGGGCACTAGGGCTCCCGGGAGACAGCAAGGCAGTAGGCTGATGATTCTTTCTTTACAGGTATTGCTTTTCCCACCAGCATTTCGGTAAATAACTGTGTATGTCACTTCTCCCCTTTGAAGAGCGACCAGGATTA
allele #1 CTACATTTCTGAAGGGCACTAGGGCTCCCGGGAGACAGCAAGGCAGTAGGCTGATGATTCTTTCTTTACAGGTATTGCTTTTCCCACCAGCATTTCGGTAAATAACTGTGTATGTCACTTCTCCCCT--------CGACCAGGATTA (Δ 8 bp)
allele #2 CTACAT-----------------------------------------------------------------------------------------------------------------------------------ACCAGGATTA (Δ 131 bp)
```

**Supplementary Figure 1.** Nucleotide sequences of the *ITAF45* gene locus in WT HEK293T cells and in the *ITAF45* KO cell line (alleles 1 and 2 shown). The second intron sequence is shown in blue, and the third exon in green. Nucleotides corresponding to the sgRNA used for CRISPR-mediated genome editing are underlined. In allele-1, an 8-bp deletion occurred, shifting the reading frame at the beginning of the third exon. Allele-2 has a 131-bp deletion disrupting the intron-exon junction and affecting pre-mRNA splicing.

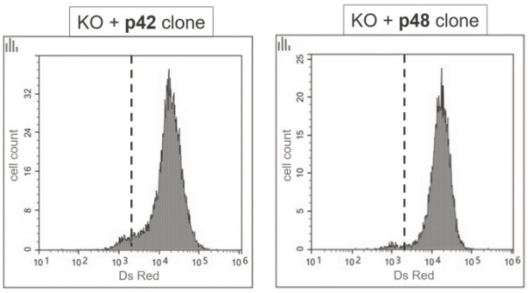


**Supplementary Figure 2.** Validation of ITAF45-KO+p42 and ITAF45-KO+p48 monoclonal cell lines for homogeneous transgene expression across the cell population. The mCherry gene present in the lentiviral cassette was used as a marker of transgene expression. Representative flow cytometry plots for the two cell lines are shown. mCherry-negative cells are to the left of the dashed line, whereas the vast majority of cells are mCherry-positive.

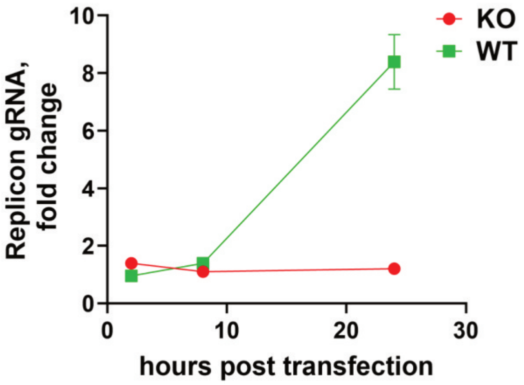


**Supplementary Figure 3.** Mengo replicon RNA accumulation in WT and KO cells transfected with in vitro-transcribed replicon RNA, measured at 2, 8, and 24 hpt by RT-qPCR. The experiments were conducted in three biological replicates, with qPCR reactions performed in four technical replicates. The means ± SD are shown.

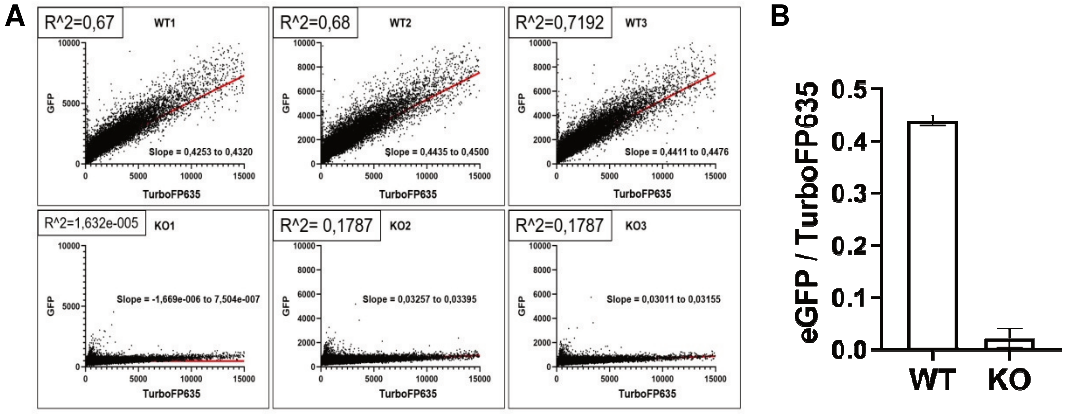


**Supplementary Figure 4.** Flow cytometry data for WT and ITAF45-KO HEK293T cells transfected with capped and polyadenylated KAT-EMCV_IRES-EGFP mRNA. (A) Flow cytometry plots from three biological replicates are shown. The analysis was performed 24 h after transfection. A linear regression was performed on the eGFP/TurboFP635 signal. $R^2$ and slope coefficients are shown. (B) Slope coefficients from three independent experiments (mean ± SD) are shown.
